# Generation of Temperature Sensitive Mutations with Error-Prone PCR in a Gene Encoding a Component of the Spindle Pole Body in Fission Yeast

**DOI:** 10.1101/595447

**Authors:** Ngang Heok Tang, Chii Shyang Fong, Hirohisa Masuda, Isabelle Jourdain, Masashi Yukawa, Takashi Toda

**Author notes:** Contributed equally (Ngang Heok Tang, Chii Shyang Fong).

## Abstract

Temperature-sensitive (ts) mutants provide powerful tools, thereby investigating cellular functions of essential genes. We report here a simple procedure to generate ts mutations using error-prone PCR in *pcp1* that encodes a spindle pole body (SPB) component in *Schizosaccharomyces pombe*. This manipulation is not restricted to analysis of Pcp1, and can be suited to any essential genes involved in other processes.

---

Microtubules (MTs) play essential roles in a variety of physiological processes, including maintenance of cell polarity, intracellular trafficking and chromosome segregation [1, 2]. One of the key events in MT assembly is its initial nucleation. The MTs are nucleated from a structural platform known as the microtubule organising centre (MTOC) [3]. The major MTOC in animal cells is the centrosome, which is made up of two perpendicular centrioles and a surrounding electron-dense pericentriolar material [4]. The centrosome equivalent in fungi is referred to as the spindle pole body (SPB), a disc-like structure associated with or embedded in the nuclear envelope [5].

In the fission yeast, *Schizosaccharomyces pombe*, the SPB resides on the cytoplasmic surface of the nuclear envelope during interphase. Upon entry into mitosis, it undergoes structural alterations, thereby being inserted into the nuclear envelope and activated to nucleate spindle MTs. Although the structures and organisation of the centrosome and the SPB are different, these two organelles have many components in common due to their functional similarity [6, 7]. These include γ-tubulin that comprises a multi-subunit complex referred to as the γ-tubulin complex (γ-TuC), receptors of the γ-TuC on the SPB (e.g. Pericentrin/Pcp1 and CDK5RAP2/Mto1) and Centrin (Cdc31) [6, 7]. The γ-TuC plays a central role in MT nucleation and assembly, in which Pcp1 is responsible to localise the γ-TuC to the nuclear side of the SPB during mitosis [8, 9, 10]

Conditional mutants provide powerful tools for functional analysis of essential genes. A temperature sensitive (ts) mutant is able to grow at the permissive temperature (e.g. 27°C) like wild-type (WT) cells and display defective phenotypes only at the restrictive temperature (e.g. 36°C). Several methodologies, including spontaneous mutations and mutagen-induced or site-directed mutagenesis have been successfully used to generate ts mutants in *S. pombe*. When we wish to create ts mutants in particular genes (gene of interest, GOI), the implementation of error-prone polymerase chain reaction (PCR) becomes instrumental; with this methodology, we can generate desired ts mutations in GOI easily and efficiently. Here, we describe screening for ts mutants using this error-prone PCR method in the *pcp1* gene, which encodes the vertebrate Pericentrin homologue.

The procedure for isolation of ts mutants in *S. pombe* is schematically shown in Fig. 1. In order to make the subsequent selection process easier, firstly, a WT strain that contains a drug resistance marker in the 3’ end of the GOI (*pcp1* in this case) was constructed with a standard procedure [11] (Fig. 1, I). Available drug resistance markers include G418 (kan^R^), hygromycin B (hph^R^) and nourseothricin (ClonNat^R^) [12]. To insert a kan^R^ cassette into the 3’ end of the *pcp1* gene, the cassette was amplified using a set of 100-mer primers in which 80 bases target the *pcp1* gene and 20 bases target the resistance cassette. Upon transformation of *S. pombe* cells with the PCR fragments [11, 12], we picked up several colonies that were formed on G418-containing plates and checked correct insertion as follows. A small amount of cells from a single colony was transferred to a PCR tube containing a freshly prepared lysis buffer (40 mM NaOH, 0.01% sarcosyl). The sample was boiled at 95°C for 15 minutes, placed on ice for 3 minutes and vortexed for 3-5 seconds. The cells were centrifuged at 5,000 rpm for 10 seconds and 1.5 μL of the supernatant was added to the 18.5 μL PCR reaction mix.

**Figure 1.**
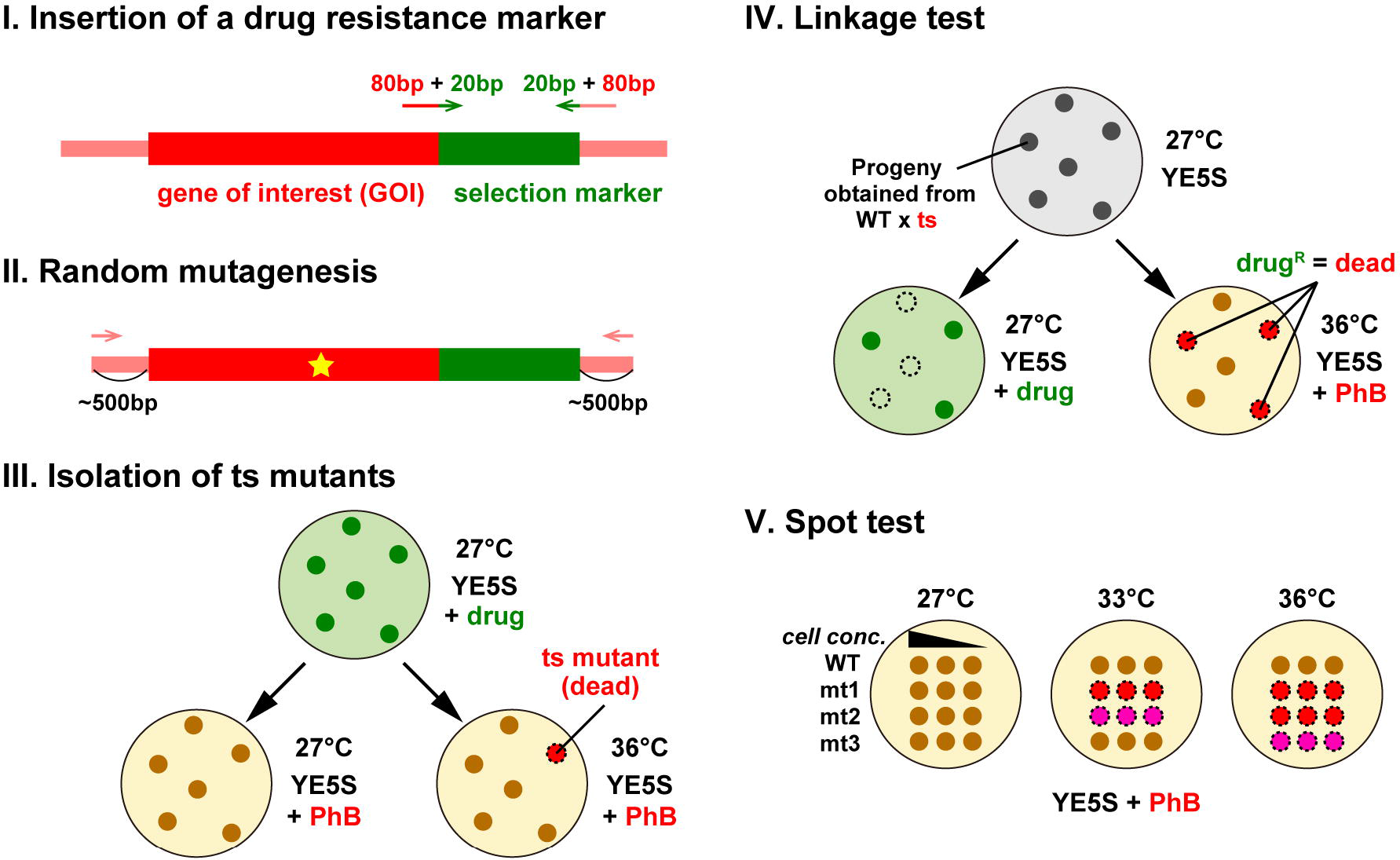
Schematic diagram illustrating the procedure for isolation of temperature sensitive mutants. Randomly mutagenised PCR fragments of GOI containing a drug resistance marker gene are transformed into a wild type fission yeast strain (I-II). Cells are allowed to grow on non-selective medium overnight for endogenous gene replacement to occur. Transformants are then selected on plates containing the selection drug at 27°C for 3 to 4 days (III). Replicated plates are incubated at 27°C and 36°C. Temperature sensitive mutants are those that do not grow at 36°C, but grow normally at 27°C. Phloxine B (PhB) is used to help detect dead cells, which are stained dark red with this dye. Candidate ts colonies are backcrossed with a wild type strain to examine co-segregation between the ts phenotype and drug resistance (IV). Spot tests are performed at various temperatures to determine the restrictive temperature (e.g. 33°C and 36°C, V).

Next, we performed random mutagenesis by error-prone PCR to introduce mutations within the *pcp1* gene (Fig. 1, II). In brief, the *pcp1* gene is amplified by PCR using unbalanced dNTPs, where one of the nucleotides is ten times more concentrated than the three others, thereby rendering the PCR amplification error-prone. Experimental conditions for amplification of PCR products may vary and need to be tested and optimised (e.g. the selection of appropriate DNA polymerases to be used). Thus far, we have obtained reproducible and reliable results with a combination of 10× dGTP and Vent DNA polymerase (New England Biolabs). It is important to use the genomic DNA derived from the strain carrying a kan^R^ marker insertion in the 3’ end of the *pcp1* gene as a template. The amplified fragment would consist of 5’ 500bp-GOI-kan^R^ cassette-500bp 3’. The presence of 500 bp in each direction facilitates homologous recombination for incorporation of the fragment into the endogenous *pcp1* locus upon transformation.

Transformants were spread on non-selective YE5S plates. After incubation at 27°C for 1 day, the cells were replica-plated onto YE5S+G418 selection plates. Upon further 3-4 days incubation at 27°C, kan^R^ colonies would appear (Fig. 1, III). These colonies were then replica-plated onto two YE5S and two YE5S+Phloxine B plates. One set of YE5S and YE5S+Phloxine B plates was incubated at 27°C and another set was placed at 36°C for 1-2 days. We visually identified candidates of ts mutants by their ability to form colonies at 27°C, but not at 36°C (Fig. 1, III). Positive ts clones should be stained dark pink at 36°C on YE5S plates containing Phloxine B, which stains dead cells. Cells of each candidate colony could be picked up from plates incubated at 36°C and observed under a benchtop microscope to assess cell morphology.

To confirm the linkage between the ts phenotype and the *pcp1* mutations, each ts candidate was backcrossed with a WT strain at 27°C, followed by random spore analysis [13]. After spores were plated onto YE5S plates and incubated at 27°C for 3 days, colonies were replica-plated onto YE5S, YE5S+G418 and two YE5S+Phloxine B plates. We incubated each of YE5S, YE5S+G418 and YE5S+Phloxine B plates at 27°C, while the other YE5S+Phloxine B plate was placed at 36°C for checking the ts phenotype. Note that a correct ts mutant should always show co-segregation between temperature sensitivity and G418 resistance (Fig. 1, IV).

To assess the ts phenotype in details, we performed spot tests. This test is a semi-quantitative growth assay where the same number of cells from different strains are “spotted” on plates, which, in this instance, are incubated at various temperatures. This assay allows us to identify restrictive (no growth), semi-permissive (weak growth) or permissive temperature (normal growth) for individual ts mutants (Fig. 1, V). In this study, we isolated seven *pcp1* mutants exhibiting growth defects at 36°C (Fig. 2a). Among these ts mutants, four mutants (*pcp1−14, −15, −16, −17* and −*18*) were sequenced; each mutant contained different point mutations within Pcp1 (Fig. 2b) [9, 10].

**Figure 2.**
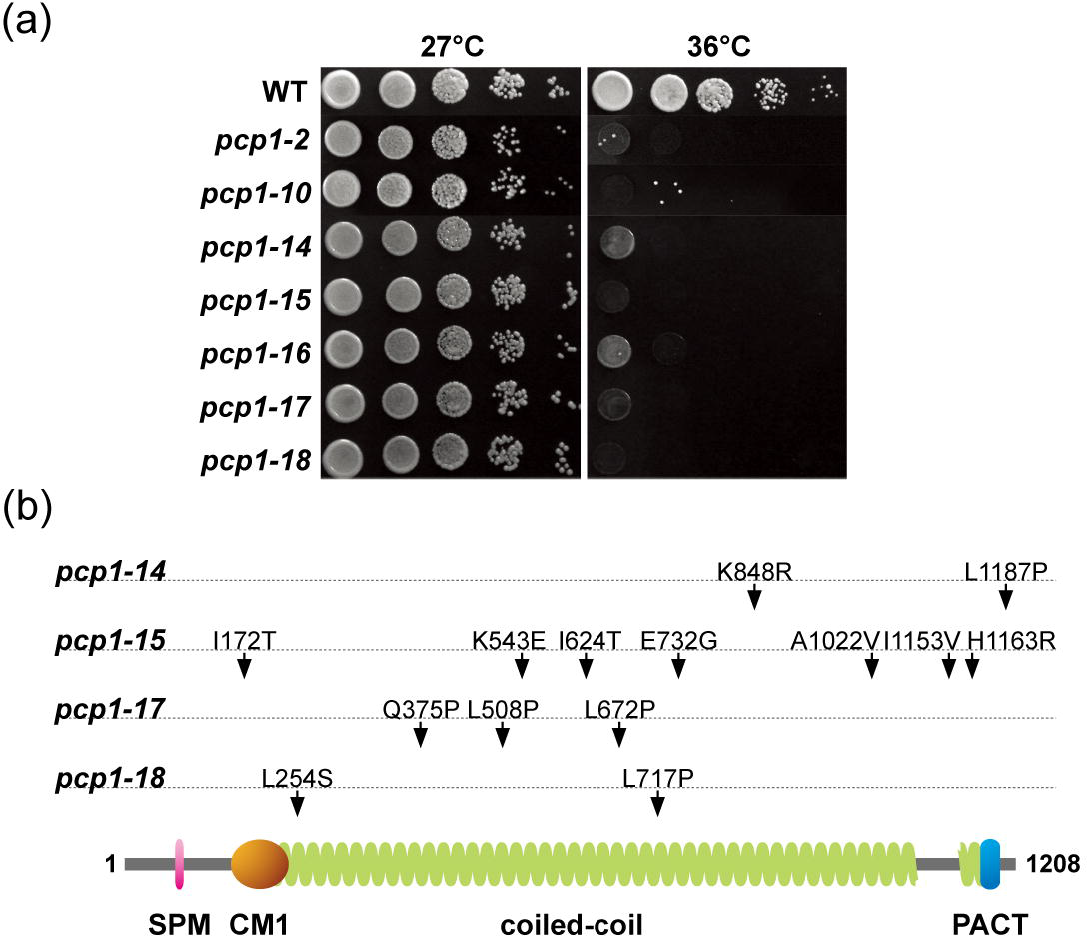
Spot tests and mutated amino acid residues within individual *pcp1* mutants. **a.** Spot test. 5 × 10^4^ cells of individual strains were spotted on the left-most line on YE5S plates and ten-fold serial dilution assays were performed. Plates were incubated at 27°C or 36°C for 3 days. **b.** Summary of mutation sites in the *pcp1* ts mutants. Functional domains within Pcp1 (e.g. SPM, CM1 and PACT) [10, 14] are indicated together with mutated amino acid residues in individual ts mutants.

The procedure presented here is applicable not only for the creation of ts mutants for a gene encoding an SPB component but also for any essential genes of interest. The strategy is based upon a simple principle, and experimentation is straightforward, by which everybody working on molecular genetics and cell biology in the fission yeast system can follow.

## Acknowledgments

We thank Dr Yoshinori Watanabe for initially implementing this method and introducing it to us.

Step-by-step protocol for isolation of ts mutants is available on line as Supplemental Material.

## Funding Acknowledgements

This work was supported by the Japan Society for the Promotion of Science KAKENHI Scientific Research (A) under Grant [16H02503]; Challenging Exploratory Research under Grant [16K14672] (T.T.); and KAKENHI Scientific Research (C) under Grant [16K07694] (M.Y.).

## Author contributions

N.H.T. and C.S.F performed the experiments and H.M. and I.J. supported them. T.T. and M.Y. wrote the manuscript with help from N.H.T., C.S.F. H.M. I.J‥

## References

1. Luders J, Stearns T. Microtubule-organizing centres: a re-evaluation. Nat Rev Mol Cell Biol. 2007 Feb;8(2):161–167. PubMed PMID: 17245416.

2. Nogales E. Structural insights into microtubule function. Annu Rev Biochem. 2000;69:277–302.

3. Bettencourt-Dias M, Glover DM. Centrosome biogenesis and function: centrosomics brings new understanding. Nat Rev Mol Cell Biol. 2007 Jun;8(6):451–463. PubMed PMID: 17505520.

4. Petry S, Vale RD. Microtubule nucleation at the centrosome and beyond. Nat Cell Biol. 2015 Aug 28;17(9):1089–1093. doi: 10.1038/ncb3220. PubMed PMID: 26316453.

5. Ding R, West RR, Morphew M, et al. The spindle pole body of *Schizosaccharomyces pombe* enters and leaves the nuclear envelope as the cell cycle proceeds. Mol Biol Cell. 1997;8:1461–1479.

6. Cavanaugh AM, Jaspersen SL. Big lessons from little yeast: budding and fission yeast centrosome structure, duplication, and function. Annu Rev Genet. 2017 Nov 27;51:361–383. doi: 10.1146/annurev-genet-120116-024733. PubMed PMID: 28934593.

7. Fu J, Hagan IM, Glover DM. The centrosome and its duplication cycle. Cold Spring Harb Perspect Biol. 2015;7(2):a015800. doi: 10.1101/cshperspect.a015800. PubMed PMID: 25646378.

8. Flory MR, Morphew M, Joseph JD, et al. Pcp1p, an Spc110p-related calmodulin target at the centrosome of the fission yeast *Schizosaccharomyces pombe*. Cell Growth Differ. 2002;13(2):47–58. PubMed PMID: 11864908.

9. Tang NH, Okada N, Fong CS, et al. Targeting Alp7/TACC to the spindle pole body is essential for mitotic spindle assembly in fission yeast. FEBS Lett. 2014 Jun 14;588(17):2814–2821. doi: 10.1016/j.febslet.2014.06.027. PubMed PMID: 24937146; Eng.

10. Fong CS, Sato M, Toda T. Fission yeast Pcp1 links polo kinase-mediated mitotic entry to γ-tubulin-dependent spindle formation. EMBO J. 2010 Jan 6;29(1):120–130. doi: 10.1038/emboj.2009.331. PubMed PMID: 19942852; PubMed Central PMCID: PMCPMC2788132. eng.

11. Bahler J, Wu JQ, Longtine MS, et al. Heterologous modules for efficient and versatile PCR-based gene targeting in *Schizosaccharomyces pombe*. Yeast. 1998 Jul;14(10):943–951. doi: 10.1002/(SICI)1097-0061(199807)14:10<943::AID-YEA292>3.0.CO;2-Y. PubMed PMID: 9717240.

12. Sato M, Dhut S, Toda T. New drug-resistant cassettes for gene disruption and epitope tagging in *Schizosaccharomyces pombe*. Yeast. 2005 May;22(7):583–591. doi: 10.1002/yea.1233. PubMed PMID: 15942936.

13. Moreno S, Klar A, Nurse P. Molecular genetic analysis of fission yeast *Schizosaccharomyces pombe*. Methods Enzymol. 1991;194:795–823. PubMed PMID: 2005825.

14. Lin TC, Neuner A, Schlosser YT, et al. Cell-cycle dependent phosphorylation of yeast pericentrin regulates γ-TuSC-mediated microtubule nucleation. eLife. 2014;3:e02208. doi: 10.7554/eLife.02208. PubMed PMID: 24842996; eng.

