## Supplementary material for "Generation of Temperature Sensitive Mutations with Error-Prone PCR in a Gene Encoding a Component of the Spindle Pole Body in Fission Yeast": Supple file

**Step-by-step protocol for isolation of temperature sensitive mutations in gene of interest (GOI)**

***1. Replica Plating***

1. Assemble the replica plating kit (**Fig. S1a**). Briefly, a piece of cling film is secured on top of a block by a rubber band. This forms the base of a replica kit (*See* **Note 1**). Then, assemble a piece of Whatman paper on top of a piece of tissue paper. Secure them onto the block using another band or a metal ring. This forms the filter paper used for replica plating.
2. Apply the plate to be replicated onto the Whatman paper. Tap gently with the palm of the hand in order to ensure all the colonies on plate are transferred to the Whatman paper (**Fig. S1b**).
3. Apply selection plates onto the replica kit containing colonies. Gently tap to ensure even distribution of the agar-paper surface (**Fig. S1b**).
4. Remove the plate from the Whatman paper (*See* **Note 2**).
5. Steps 3 & 4 can be repeated for multiple times with other selection plates, if desired.

***2. Construction of a strain containing a drug resistance marker in the 3' end of the GOI***

1. Set up a PCR reaction to amplify a drug resistance marker fragment (**Fig. S2a, b**).
2. Run 1-2 μL of the amplified PCR fragment on 1% agarose gel. For the kan$^R$ cassette, the correct PCR fragment should be of ~1.6 kb long.
3. Concentrate the remaining PCR fragments into 10 μL of double-distilled water (*See* **Note 3**).

***3. Transformation of PCR fragments into S. pombe cells***

1. Before starting, boil the salmon sperm DNA for 5 min, then leave on ice (2 μL of a 10 mg/mL stock will be needed per transformation).
2. Harvest 50 mL exponentially growing cells by centrifugation at 3000 rpm for 3-5 minutes.
3. Discard supernatant and wash the cells with 10 mL of Lithium Acetate with TE (LiOAc/TE), for 5 minutes with gentle rotation. Spin them down as before. Repeat this step once.

4. Resuspend the cells in 100 μL of LiOAc/TE and transfer the cell solution into a 1.5 mL microcentrifuge tube.
5. Add 7.5 μL of carrier DNA and 10 μL DNA fragments into the cell solution (*See* **Note 4**).
6. Add 260 μL of 40% PEG/LiOAc/TE and vortex briefly for 3 seconds.
7. Incubate the solution at 27˚C for 2 hours, shaking at 220 rpm.
8. Add 43 μL of DMSO and vortex briefly for 3 seconds.
9. Heat shock at 42˚C for 5 minutes.
10. Wash the cells with 500 μL of YE5S. Spin down at 5000 rpm for 10 seconds.
11. Discard supernatant and resuspend the cells in 500 μL of YE5S.
12. Incubate the cells at 27˚C for 1.5 hours, shaking at 220 rpm (*See* **Note 5**).
13. Spin down the cells at 5000 rpm for 10 seconds. Discard 200 μL of the supernatant and resuspend the cells in the remaining 300 μL of YE5S.
14. Plate 150 μL of cell solution onto YE5S plate, using glass balls. Shake gently to evenly spread the solution on the plate (*See* **Note 6**).
15. Replica-duplicate the colonies from these YE5S plates into YE5S+G418 plates. Incubate plates at 27˚C for 4 to 5 days.
16. Cells that have grown on G418 plates should have kan$^{R}$ cassette inserted into the GOI. To confirm this, check correct insertion of the kan$^{R}$ cassette by colony PCR.

### *4. Colony PCR*

1. Pick a small amount of cells from a single colony and transfer to a PCR tube containing solution.
2. Suspend the cells in 12 μL of freshly prepared 40 mM NaOH/0.01% sarcosyl mix.
3. Boil the sample at 95°C for 15 minutes.
4. Place the tube on ice to cool for 3 minutes.
5. Vortex for 3-5 seconds to mix the solution.
6. Spin down the cells at 5000 rpm for 10 seconds.
7. Use 1.5 μL of the supernatant and 18.5 μL of reaction mixture for PCR (**Fig. S2c, d**).

### *5. Random mutagenesis by error-prone PCR*

1. Purify genomic DNA from a strain containing GOI-kan$^{R}$ using the commercial MasterPure Yeast DNA purification kit (Cambio Ltd.).

2. Design a set of primers (20-30 mer) that anneal to 500 bp upstream (forward primer; 500 bp before the START codon) and 500 bp downstream (reverse primer in reverse complimentary form; 500 bp after the STOP codon) of the GOI-kan$^R$ (*See* **Note 7**).
3. Amplify the GOI-kan$^R$ fragment with error-prone PCR using Vent DNA polymerase (New England Biolabs) supplemented with 10× dGTP (100 mM for dGTP; 10 mM for dATP, dTTP and dCTP).
4. Run a portion (1-2 μL is sufficient) of the amplified PCR fragment on 1% agarose gel.
5. If a product was amplified at the right size and in sufficient amount (a total of approx. 400 μg are needed for each PCR), re-do another 40 PCRs using the same conditions.
6. Using a commercial kit or by ethanol precipitation, concentrate each PCR reaction individually into 10 μL of water.
7. Transform each concentrated PCR reaction twice into WT *S. pombe* cells (i.e. ideally 80 transformations) (*See* **Note 8**).
8. Allow cells to grow at 27$^{\circ}$C overnight for endogenous gene replacement to occur.
9. Replica the lawn of *S. pombe* cells on YE5S+G418 plates and incubate them at 27$^{\circ}$C for 3 to 4 days.

***6. Isolation of temperature sensitive (ts) mutants***

1. Replica each plate onto two YE5S and two YE5S+Phloxine B plates.
2. Incubate one YE5S and one YE5S+Phloxine B plate at 27$^{\circ}$C for 2 days. Incubate the other YE5S and YE5S+Phloxine B plates at 36$^{\circ}$C for 1 to 2 days.
3. Visually identify candidate ts mutants by their ability to grow and form a colony at 27$^{\circ}$C, but not at 36$^{\circ}$C. Positive clones should not only grow less at 36$^{\circ}$C than 27$^{\circ}$C, but should also be stained pink by Phloxine B, which stains dead cells dark red (e.g. at 36$^{\circ}$C).
4. Pick some cells from each candidate colony, transfer on a microscope slide with a bit (~2 μL) of water, cover with a coverslip and observe under a benchtop microscope to rapidly assess the phenotype. Select clones that show a phenotype of interest, clones that have different phenotypes, and clones that have a range of temperature sensitivity (variations of red colours on Phloxine B plates).
5. Re-streak candidate ts mutants from the YE5S plates incubated at 27$^{\circ}$C, onto fresh YE5S plates. Incubate at the permissive temperature 27$^{\circ}$C.

***7. Backcrossing of ts mutants and random spore analysis***

1. Cross each ts candidate with a WT strain of the opposite mating type.
2. Inoculate a loopful of asci mix (about one colony size) into 100 µL of 0.5~1% glusulase to breakdown the ascus wall. Incubate tubes at 27°C for 2 hours.
3. Add 43 µL of ethanol 95% to the tube to kill any non-sporulating cells. Incubate tubes at room temperature for 30 minutes.
4. Pellet the spores by centrifugation (2000 rpm, 1 minute) and resuspend them in 1 mL of $H_2O$.
5. Plate onto YE5S plates (2 µL, 10 µL or 50 µL depending on the efficiency of sporulation) and incubate at 27°C for 3 days.
6. Replica each plate onto 1× YE5S, 1× YE5S+G418 and 2× YE5S+Phloxine B plates.
7. Incubate the YE5S, YE5S+G418 and 1× YE5S+Phloxine B at 27°C; and the other YE5S+Phloxine B at 36°C.
8. Check the plates after 2 days. A correct ts mutant should always show co-segregation between temperature sensitivity and G418 resistance.

***8. Assessment of temperature sensitivity by spot test***

1. Propagate exponentially growing cultures of WT and ts mutants in YE5S broth.
2. Determine cell density with a cell counter.
3. Dilute cultures to $2 \times 10^7$ cells/mL and transfer 100 µL of the diluted culture to a 96-well plate.
4. Prepare four ten-fold serial dilutions in the 96-well plate.
5. Spot cells onto YE5S plates with a 48-pin replicator.
6. Incubate plates at temperatures ranging from 22°C to 36°C, including one at 27°C, for 1 to 3 days.

***9. Notes***

1. This base (wooden block + cling film) does not have to be changed for every replica plating. The papers have to be changed when a set of replica plating is done.
2. Colonies will be transferred from the Whatman paper to the plate. A "colony shape" may be seen on the plate, which contains many cells. However, no colony will be visible on the plate at this point.

3. We use QIAquick PCR purification kit (Qiagen) for this. Elute the PCR fragments using 10 μL of double-distilled water.
4. Carrier DNA increases the transformation efficiency. We use salmon sperm DNA (Invitrogen). 250 ng of DNA fragments are usually sufficient per tube.
5. This step will enhance recovery of the cells.
6. We use 2 plates for each transformation, to avoid cells over-growing in a single plate. This also creates a back-up in the case of contamination. Direct plating could be performed onto YE5S+G418 plates. However, plating onto YE5S plates will enhance recovery of the cells.
7. Primers need not to be long as the amplified, mutated fragments will itself recombine with the endogenous WT gene. The amplified fragment should contain 500bp-GOI-kan$^R$ cassette-500bp. Since the kan$^R$ cassette was inserted immediately downstream of the STOP codon. The presence of 500 bp in each direction facilitates homologous recombination for incorporation of the fragment into the genomic locus after transformation. It is not a problem to subject the kan$^R$ fragment for error-prone PCR. If the cassette is mutated in error-prone PCR, it should not grow in YE5S+G418 plate during the selection process.
8. Each PCR tube may have produced and amplified different point mutations. In order not to select several times the same mutant, it is preferable not to pull all PCR products together but to transform them individually.

Figure S1. Tang et al.

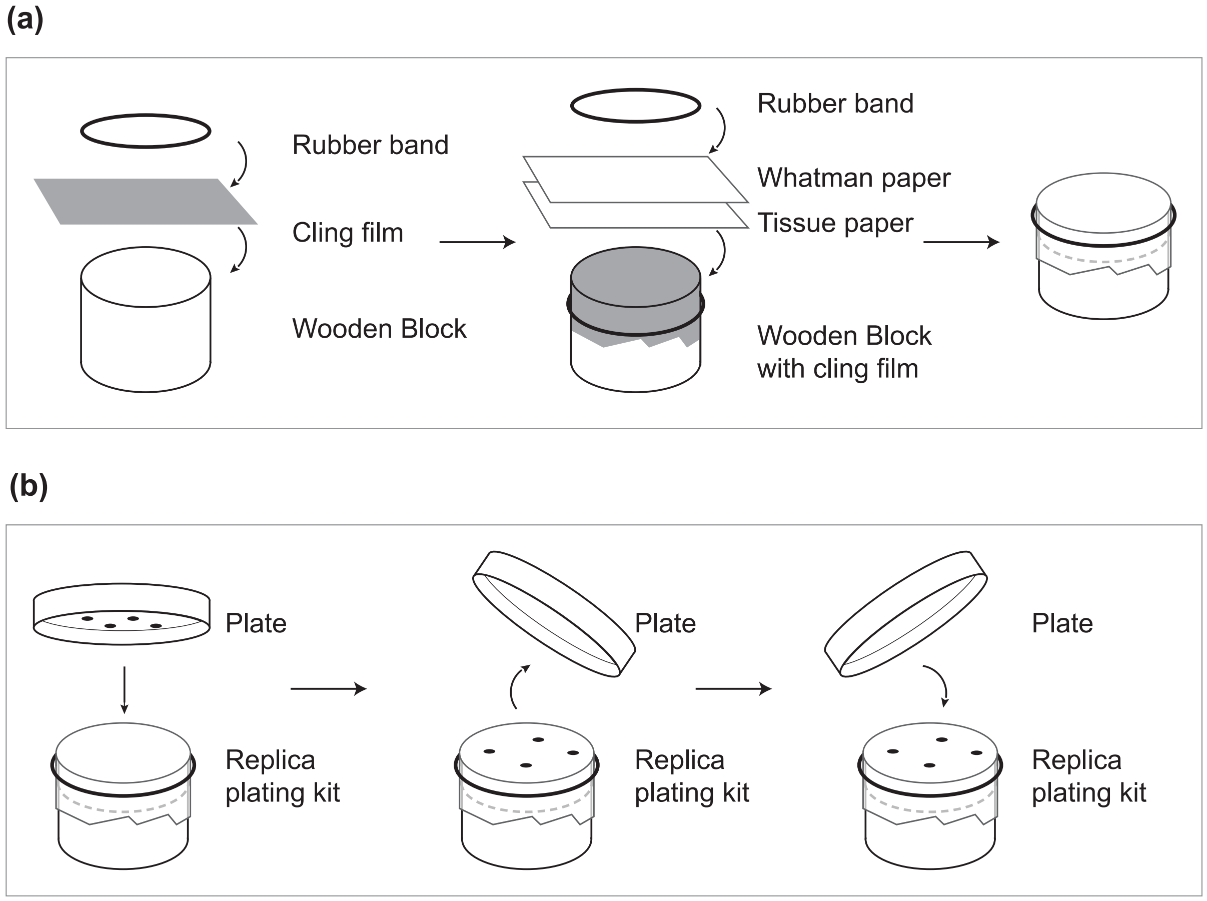


**Figure S1. Replica plating**

**(a)** Assembly of the replica plating kit. **(b)** Replica plating of plates. See Section 1 of the supplemental material for the detailed procedures.

Figure S2. Tang et al.

**(a)**

| | |
|---|---|
| 5X PrimeSTAR Buffer ($Mg^{2+}$ Plus) | 10 µl |
| dNTP Mixture (2.5 mM each) | 4 µl |
| Forward Primer | 50 pmol |
| Reverse Primer | 50 pmol |
| Template (pFA6a-kanMX6) | 5-10 ng |
| PrimeSTAR HS DNA Polymerase (2.5 U/µl) | 0.5 µl |
| Double-Distilled Water | up to 50 µl |

**(b)**

| | | |
|---|---|---|
| 94˚C | - 1 min | |
| 98˚C | - 10 sec | 35 cycles |
| 52˚C | - 15 sec | 35 cycles |
| 72˚C | - 2 min | 35 cycles |
| 72˚C | - 2 min | |
| 4˚C | - storage | |

**(c)**

| | |
|---|---|
| 10X Z buffer | 10 µl |
| dNTP Mixture (2.5 mM each) | 4 µl |
| Forward Primer | 10 pmol |
| Reverse Primer | 10 pmol |
| Template (supernatant) | 1.5 µl |
| Z taq | 0.5 µl |
| Double-Distilled Water | up to 20 µl |

**(d)**

| | | |
|---|---|---|
| 95˚C | - 5 min | |
| 98˚C | - 5 sec | 40 cycles |
| 55˚C | - 10 sec | 40 cycles |
| 72˚C | - 10 sec/kb | 40 cycles |
| 72˚C | - 5 min | |
| 4˚C | - storage | |

**Figure S2. Composition of PCR reaction mixture**

**(a, c)** Components of PCR reaction mixture for amplification of a drug-resistance cassette **(a)** and for colony PCR **(c)**, as indicated by manufacturer's instructions (#R006A, Takara). **(b, d)** Conditions used for PCR reaction.
